## Supplementary Material for "Downregulation of Dickkopf-3, a Wnt antagonist elevated in Alzheimer’s disease, restores synapse integrity and memory in a disease mouse model"

### SUPPLEMENTARY FIGURES AND TABLES

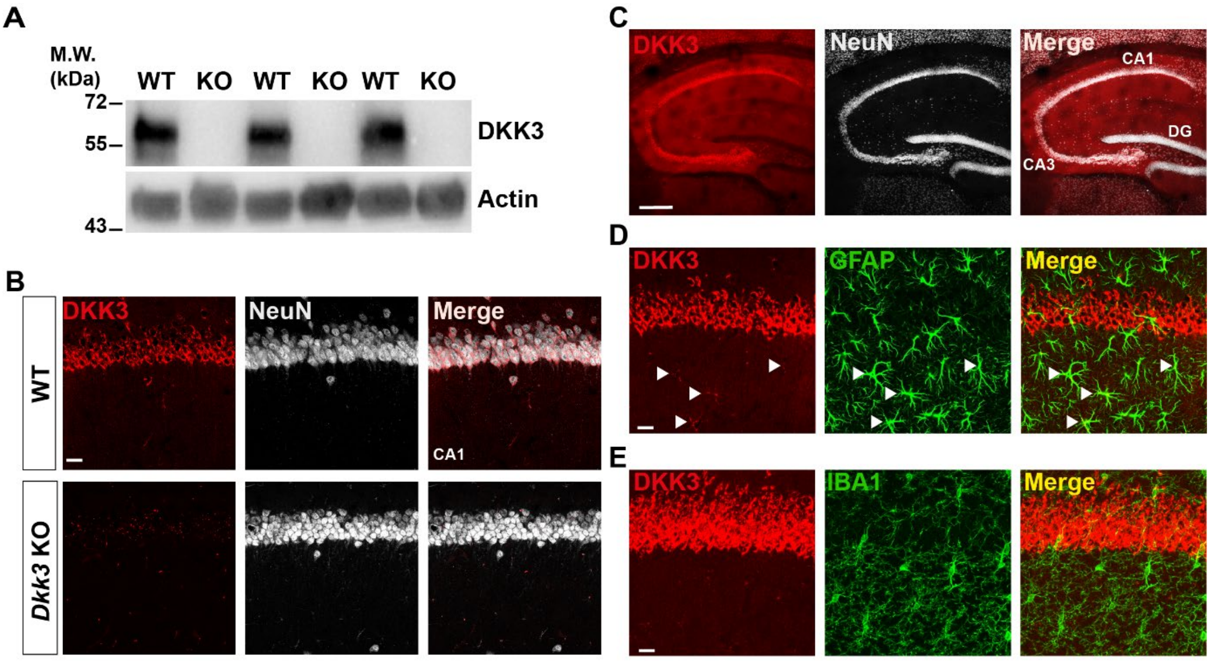


Supplementary Figure 1

**Figure S1. DKK3 is present in principal neurons of the mouse hippocampus.**

(**A**) Immunoblotting images show the presence or absence of DKK3 protein in hippocampal homogenates of WT and *Dkk3*$^{-/-}$*ApoE*$^{-/-}$ (KO) mice demonstrate the specificity of the antibody against DKK3. Actin was used a loading control.

(**B**) Confocal images show DKK3 protein in the CA1 region of the hippocampus in WT mice but absent in the *Dkk3*$^{-/-}$*ApoE*$^{-/-}$ (KO) mice. Scale bar = 25 μm.

(**C**) Confocal images of DKK3 (red) and neuronal marker NeuN (grey) demonstrate that DKK3 colocalizes with principal neurons but not with granular neurons in the hippocampus. Scale bar = 250 μm.

(**D**) Confocal images of DKK3 (red) and the astrocyte marker GFAP (green). Arrowheads indicate some astrocytes expressing low levels of DKK3. Scale bar = 25 μm.

(**E**) Confocal images of DKK3 (red) and microglial marker IBA1 (green). Scale bar = 25 μm.

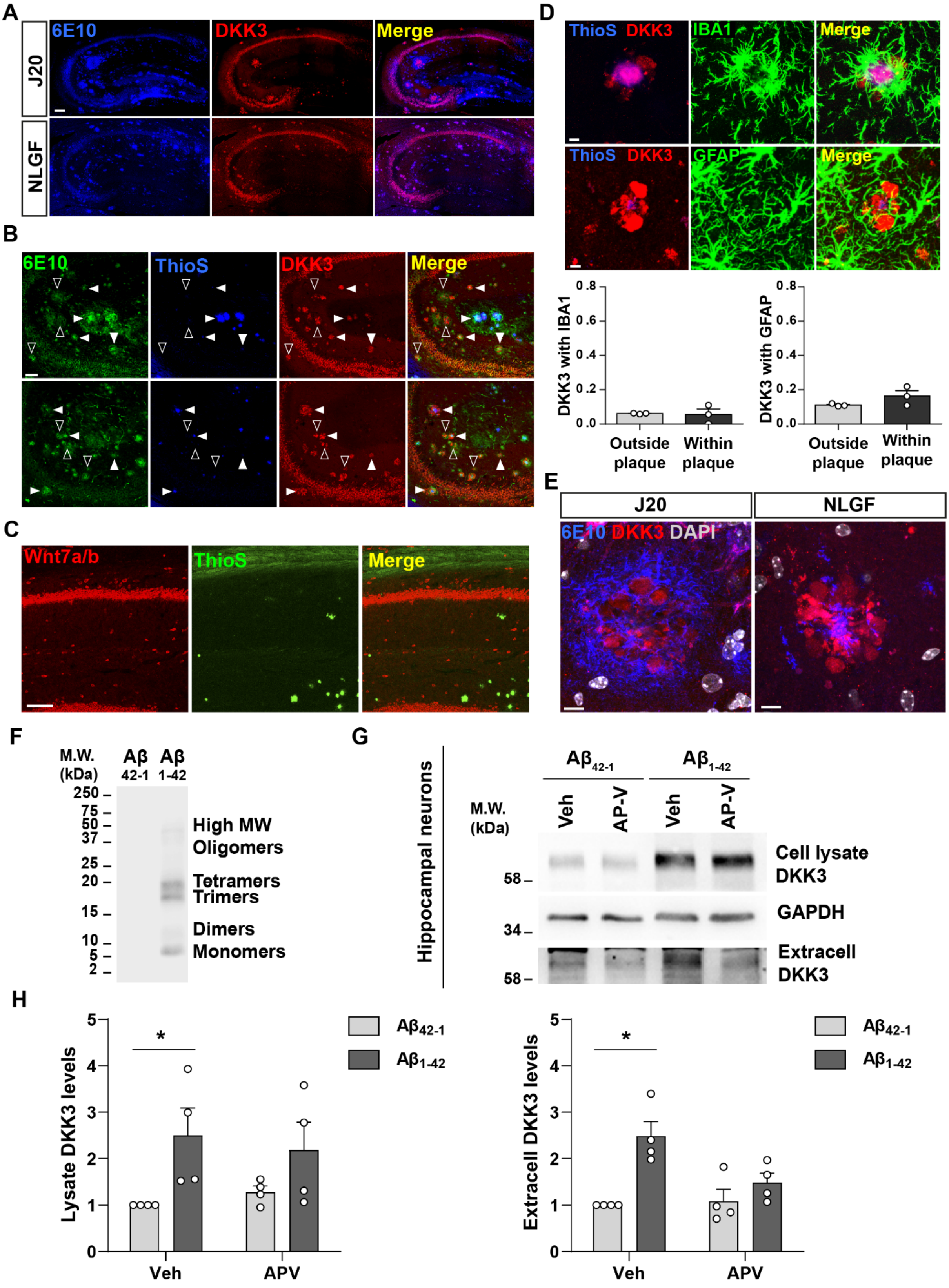


Supplementary Figure 2

**Figure S2. DKK3 accumulates in Aβ plaques in the hippocampus of J20 and NLGF mice and DKK3 levels are increased by Aβ.**

(**A**) Confocal images of Aβ plaques (6E10, blue) and DKK3 (red) in the hippocampus of 12-month-old J20 and 8-month-old NLGF mice. Scale bar = 140 µm.

(**B**) Confocal images show that DKK3 (red) accumulates in Aβ plaques (6E10, green) with or without a dense-core (ThioS, blue) in 18-month-old J20 hippocampi. Empty arrowheads indicate diffuse plaques (6E10 positive, ThioS negative) containing DKK3. White arrowheads point to dense-core plaques (positive for 6E10 and ThioS) containing DKK3. Scale bar = 58 µm.

(**C**) Confocal images of Wnt7a/b (red) and Aβ plaques stained by Thioflavin S (ThioS, green) in the CA1 region of 18-month-old J20 mice. Scale bar = 100 µm.

(**D**) Z-stack confocal images showing DKK3 (red) in Aβ plaques (ThioS, blue) and microglia (IBA1, green) or astrocytes (GFAP, green). Scale bar = 6 µm. Graphs show the Pearson's coefficient quantification showing the degree of colocalization between DKK3 and IBA1 or GFAP, n = 3 animals.

(**E**) Confocal images of Aβ plaques (6E10, blue), DKK3 (red), and DAPI staining of nuclei (grey) in the hippocampus of 12-month-old J20 and 8-month-old NLGF mice. Scale bar = 8 µm.

(**F**) Preparation of oligomers of $A\beta_{1-42}$ (Aβo). Representative immunoblots of 6E10 for $A\beta_{42-1}$ (reverse peptide control) and $A\beta_{1-42}$.

(**G**) Representative immunoblot image shows DKK3 levels in the lysate and in extracellular media of cultured hippocampal neurons treated with reverse Aβ peptide ($A\beta_{42-1}$) or Aβo ($A\beta_{1-42}$) and vehicle (Veh) or APV. GAPDH was used as a loading control in cell lysates.

(**H**) Graphs show densiometric quantifications of lysate and extracellular (extracell) DKK3 levels relative to control (Kruskal-Wallis followed by Dunn's post-hoc test; n = 4 independent experiments).

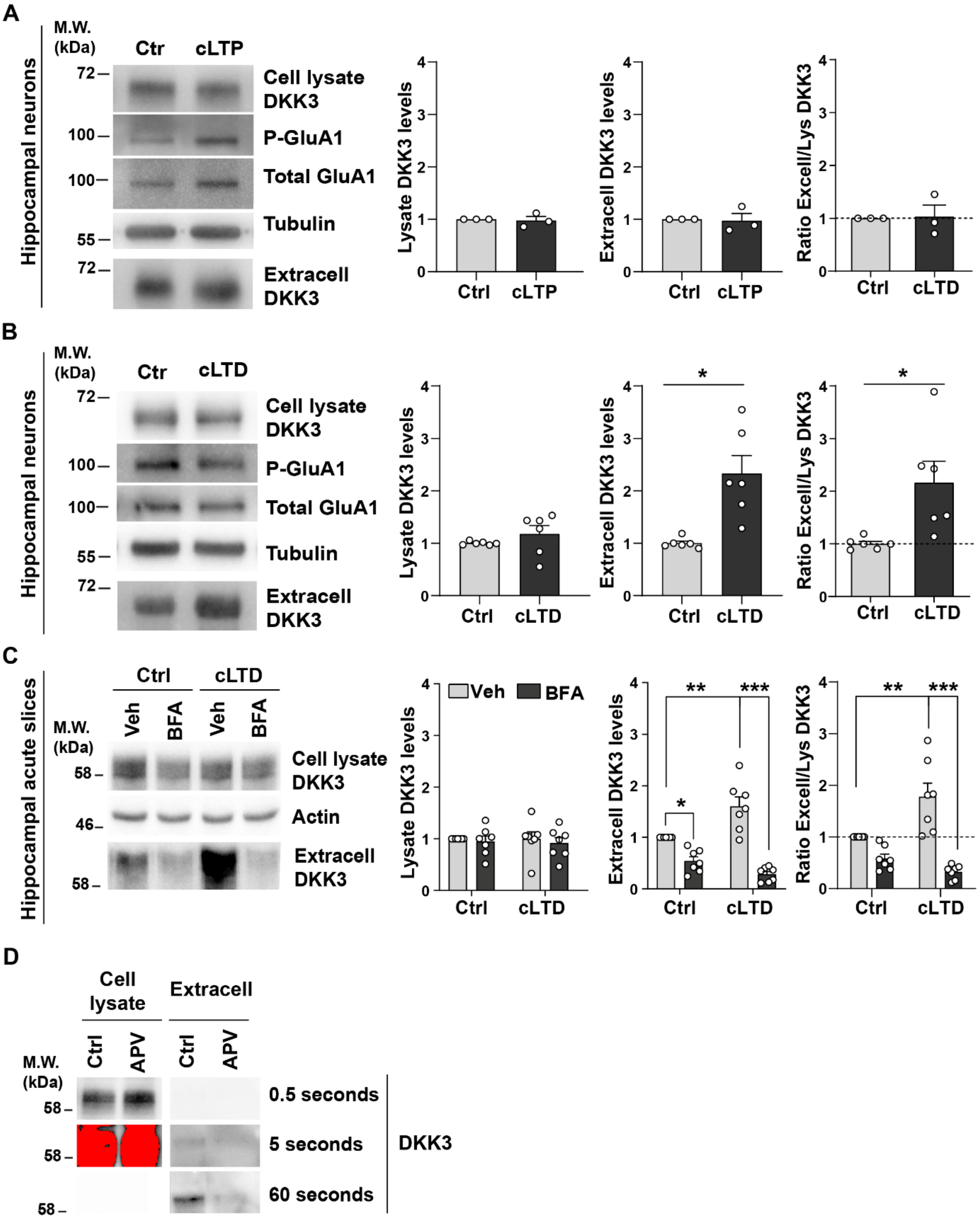


Supplementary Figure 3

**Figure S3. cLTD, but not cLTP, increases the levels of extracellular DKK3.**

(**A, B**) Hippocampal neurons were subjected to (**A**) chemical LTP (cLTP) or (**B**) chemical LTD (cLTD). DKK3 levels were analyzed in the supernatant (extracell) and cell lysates by western blot. Phospho-GluA1 Ser845 was used as a readout of LTP and LTD induction, and tubulin as a loading control for the cell lysate. Graphs show the levels of DKK3 relative to control and the ratio of extracellular/lysate DKK3 levels (Mann-Whitney Test, n = 3 independent cultures for cLTP and Student's T-test, n = 6 independent cultures for cLTD).

(**C**) Representative immunoblots show DKK3 levels in the cell lysate and extracellular (extracell) fraction of acute WT hippocampal slices treated with vehicle (Ctrl) or NMDA (cLTD) and/or Brefeldin A (BFA) for 60 minutes. Actin was used as a loading control in homogenates. Graphs show densiometric quantifications of DKK3 relative to the control condition and the ratio of extracellular/lysate DKK3 levels (Kruskal-Wallis followed by Dunn's multiple comparisons; n = 5 animals).

(**D**) Immunoblot showing DKK3 is less abundant in the extracellular fraction when compared to the cell lysate fraction. Representatives immunoblot of J20 brain slices treated with control or APV for 3 hours. Time exposure for obtaining DKK3 chemiluminescent images is indicated.

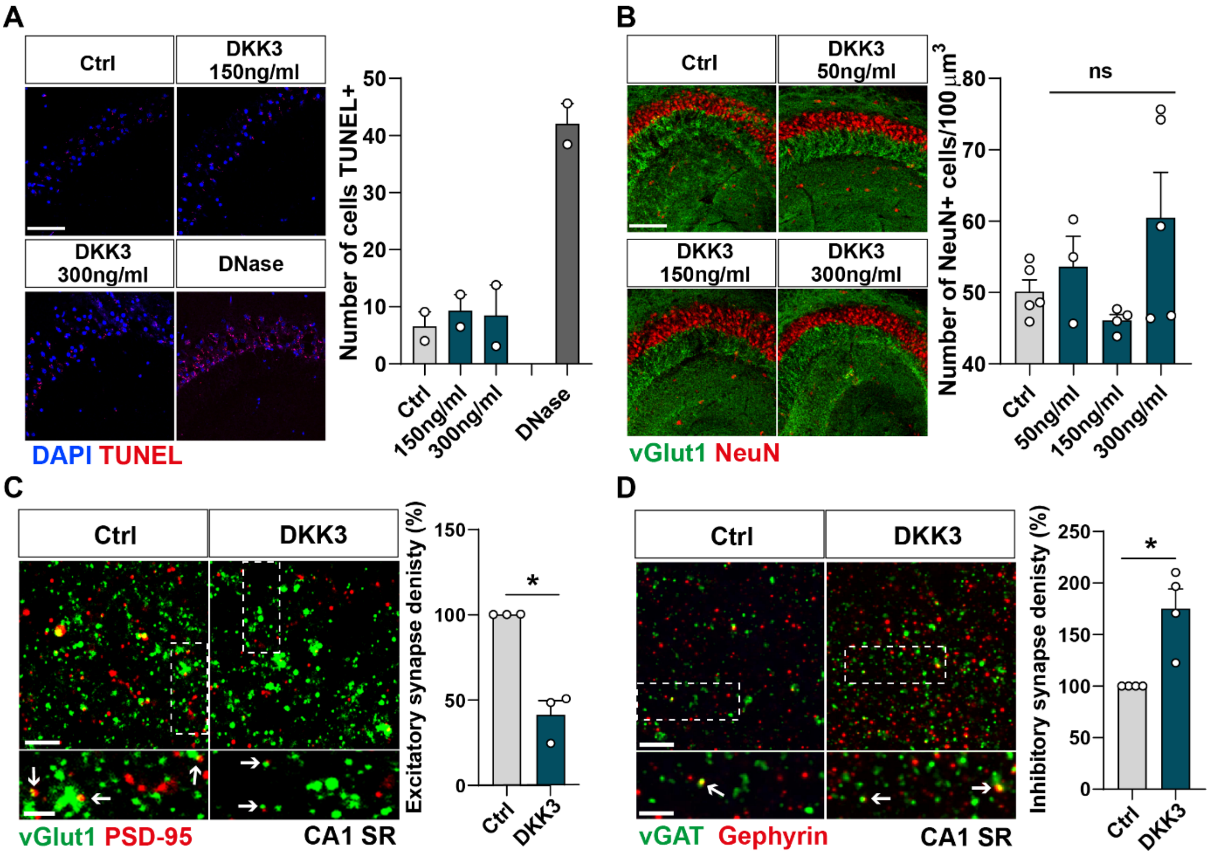


**Supplementary Figure 4**

**Figure S4. DKK3 triggers changes in excitatory and inhibitory synapses in the absence of cell death.**

(**A**) Confocal images of the TUNEL assay (red) in the hippocampus CA3 area (DAPI in blue). Graph shows the number of cells positive for TUNEL. Scale bar = 20 μm (n = 2 animals).

(**B**) Confocal images show the impact of different DKK3 concentrations on cell number (NeuN in red) and vGLUT1 (green) in the hippocampus CA3 area. Graph shows the number of NeuN+ cells per 10 0μm$^3$. Scale bar = 150 μm (One-way ANOVA followed by Tukey's post-hoc test, ns, n = 2 animals, 2-3 brain slices per animal).

(**C**) Confocal images from hippocampal CA1 SR show the effect of DKK3 on excitatory synapses (colocalized vGLUT1 puncta in green and PSD-95 puncta in red). Arrows indicate excitatory synapses. Scale bar = 5 μm and 2.5 μm in zoomed-in pictures. Quantification is shown on the right-hand side (Mann-Whitney test, n = 3 animals).

(**D**) Confocal images from hippocampal CA1 SR show the effect of DKK3 on inhibitory synapses (colocalized vGAT puncta in green and gephyrin puncta in red). Arrows point to inhibitory synapses. Scale bar = 5 μm and 2.5 μm in zoomed-in pictures. Quantification is shown on the right-hand side (Mann-Whitney test, n = 4 animals).

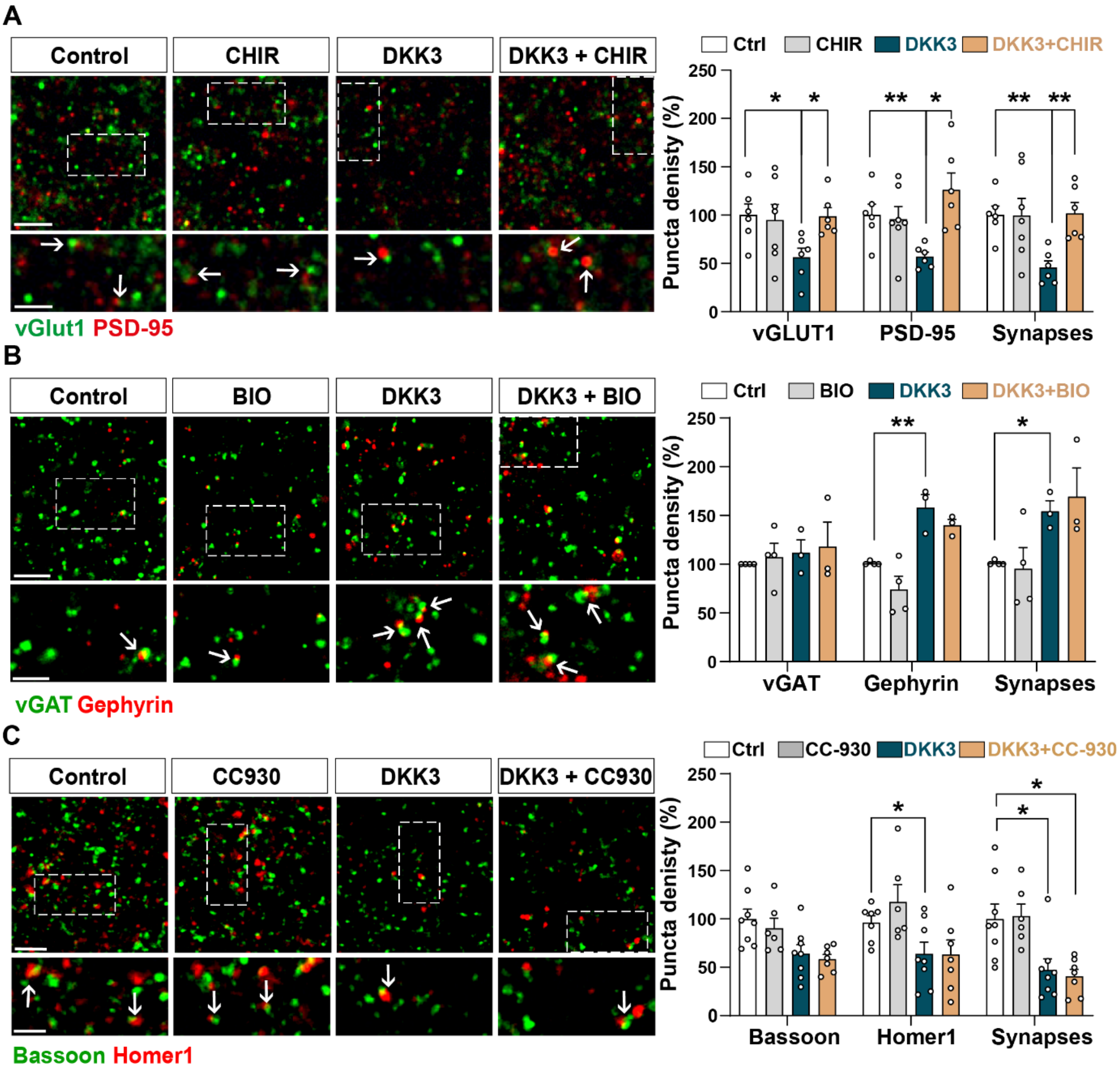


**Supplementary Figure 5**

**Figure S5. Activation of the Wnt/GSK3β pathway blocks DKK3-induced excitatory synapse loss whereas inhibition of the Wnt/JNK pathway blocks the effect of DKK3 on inhibitory synapses.**

(**A**) Confocal images show excitatory synapses (co-localized vGLUT1 in green and PSD-95 in red) in the CA3 SR after treatment with vehicle (Ctrl) or DKK3 in the absence or presence of CHIR99021 (CHIR) for 4h. Scale bar = 5 μm and 2.5 μm Graph shows the quantification of puncta density for pre and postsynaptic markers as well as excitatory synapse number as a percentage of control (Kruskal-Wallis followed by Dunn's multiple comparisons, n = 2-3 brain slices from 3 animals).

(**B**) Confocal images showing inhibitory synapses defined by the colocalization of vGAT (green) and Gephyrin (red) in the CA3 SR after treatment with vehicle (Ctrl) or DKK3 in the absence or presence of BIO. Scale bar = 5 μm and 2.5 μm. Graph shows the quantification of puncta density of pre and postsynaptic markers as well as inhibitory synapse number as a percentage of control (Kruskal-Wallis followed by Dunn's multiple comparisons, n = 3-4 animals).

(**C**) Confocal images showing excitatory synapses (colocalized Bassoon in green and Homer1 in red) in the CA3 SR after vehicle (Ctrl) or DKK3 treatment in the absence or presence of CC-930. Scale bar = 5 μm and 2.5 μm. Graph shows the quantification of pre and postsynaptic markers as well as excitatory synapse number as a percentage of control (Two-Way ANOVA followed by Tukey's multiple comparisons, n = 2-3 brain slices from 3 animals).

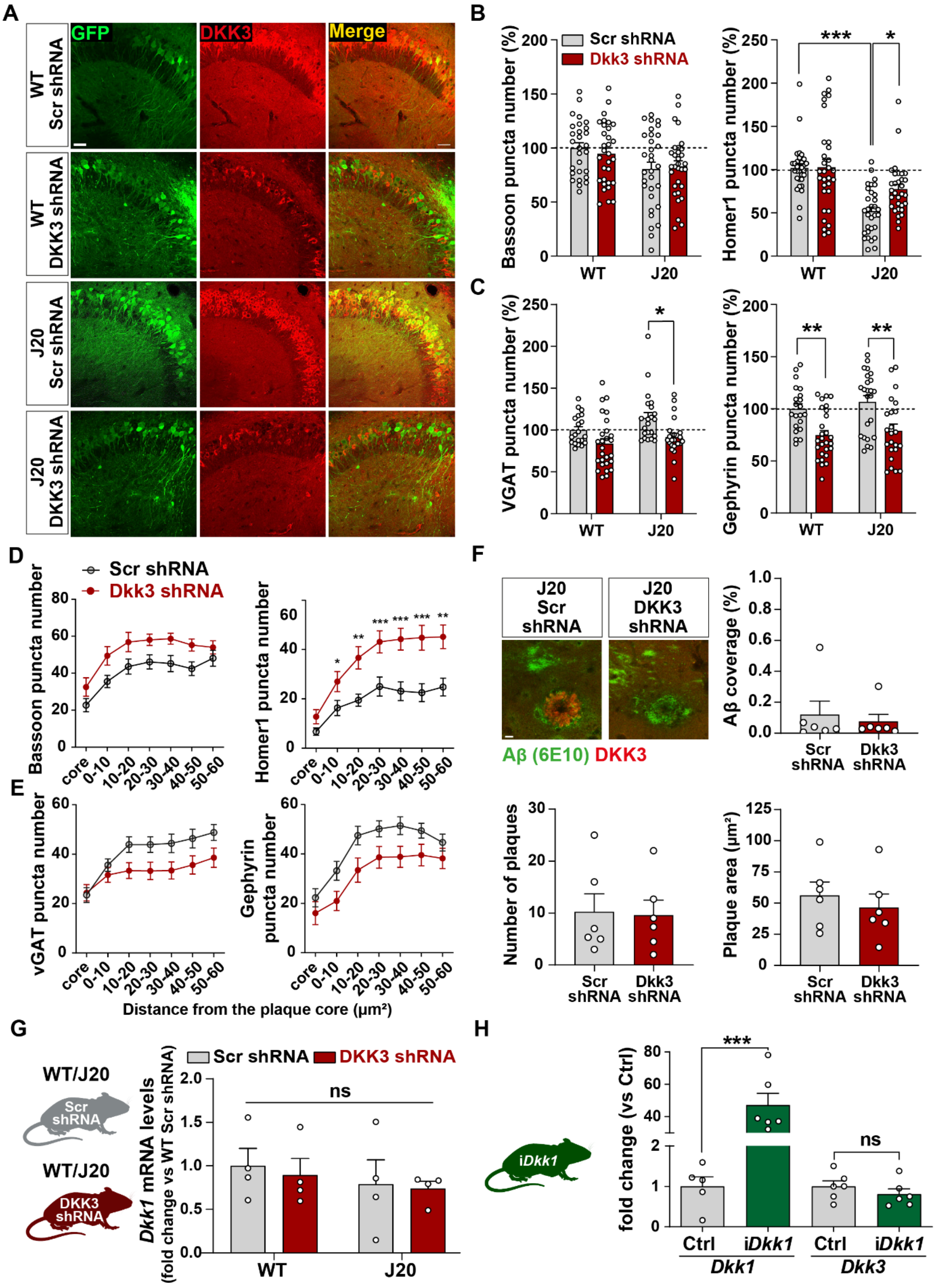


**Supplementary Figure 6**

**Figure S6. *In vivo* DKK3 loss-of-function affects pre- and postsynaptic markers but does not affect Aβ plaque pathology or *Dkk1* expression in the mouse hippocampus.**

(**A**) Confocal images of the CA3 area of the hippocampus show that endogenous DKK3 is downregulated in the hippocampus of wildtype and J20 injected with DKK3 shRNA AAV9 virus. Scale bar = 38 μm.

(**B, C**) Graphs show (**B**) excitatory (Bassoon and Homer1) and (**C**) inhibitory (vGAT and Gephyrin) puncta density as a percentage relative to WT Scr shRNA group in 4-months old WT and J20 mice (Two-Way ANOVA followed by Tukey's post-hoc test, n = 9-11 animals per condition and 3 brain slices per animal).

(**D, E**) Graphs show (**D**) excitatory (Bassoon and Homer1) and (**E**) inhibitory (vGAT and Gephyrin) puncta density at each distance from the core plaque as density per 200 $\mu m^3$ in 9-months old J20 mice (Two-Way ANOVA followed by Tukey's post-hoc test. n = 7-8 animals per condition and 3 brain slices per animal).

(**F**) DKK3 downregulation does not affect plaque load in the hippocampus. Confocal images of Aβ (6E10 in green) and DKK3 (red) in the CA3 SR of 9-months old J20 mice. Scale bar = 10 μm. Graphs show the quantification of Aβ coverage, plaque number and plaque area in the CA3 (Student's T-test, ns, n = 6 per condition).

(**G**) DKK3 downregulation does not affect *Dkk1* expression in the hippocampus. *Dkk1* expression was evaluated in the hippocampus of 4-month-old WT and J20 mice injected with Scr or DKK3 shRNA. Graph shows *Dkk1* mRNA levels normalized to the control group (Kruskal-Wallis followed by Dunn's multiple comparisons, ns, n = 4 animals per condition).

(**H**) Increased expression of *Dkk1* does not affect *Dkk3* mRNA levels. *Dkk3* expression was examined in the hippocampus of *iDkk1* mice compared to control mice after induction of *Dkk1* for 14 days. Graph shows *Dkk1* and *Dkk3* mRNA levels normalized to the control group (Student's T-test, n = 5-6 animals per condition).

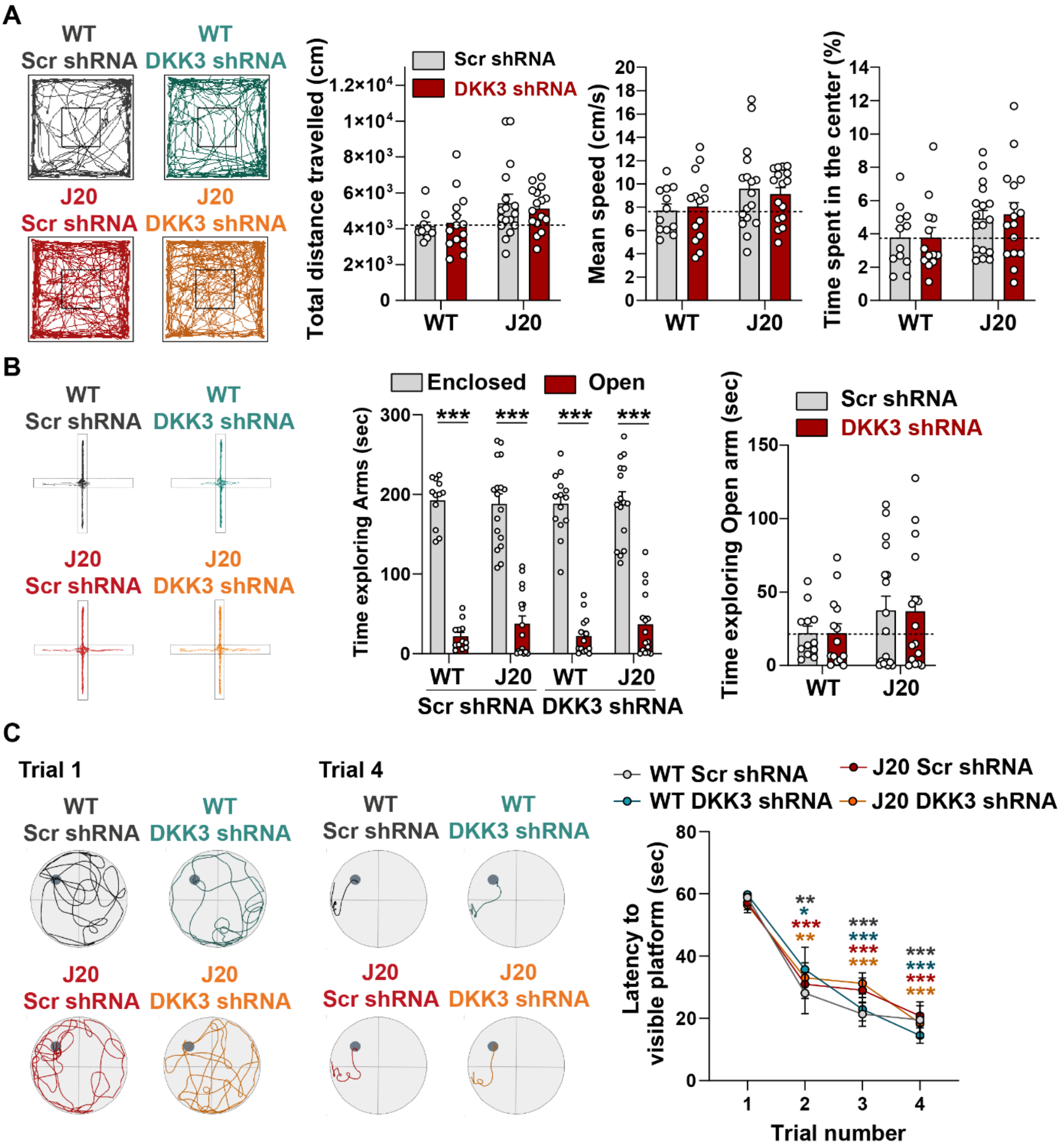


Supplementary Figure 7

**Figure S7.** ***In vivo*** **DKK3 loss-of-function does not affect locomotion or anxiety.**

(**A**) Open-field task. Representative traces of the open-field. Graphs show the total distance traveled in the open-filed (cm, left, Kruskal-Wallis test), mean speed (cm/s, middle, Two-way ANOVA) and the time spent in the center relative to the total time (%, right, Kruskal-Wallis test).

(**B**) Elevated plus-maze (EPM). Representative traces of the EPM. Left graph shows the total time exploring arms (sec, Two-way ANOVA followed by Tukey's multiple comparison test). Right graph displays the time spent in the open arms (sec, Kruskal-Wallis test).

(**C**) Morris Water Maze (MWM). Representative traces for Trials 1 and 4 in the training phase of the visible platform version of the MWM. The time (sec) to reach the hidden platform was measured and compared over trials. No differences were observed between groups. (Two-Way ANOVA followed by Tukey's post-hoc test).

**Table S1. Human brain samples information.** M = Male, F = Female, PMI = Post-mortem interval.

| Patient | Braak stages | Sex (F,M) | Age (years) |
|---|---|---|---|
| 1 | 0 | F | 85 |
| 2 | 0 | F | 54 |
| 3 | 0 | F | 72 |
| 4 | 0 | M | 81 |
| 5 | 0 | F | 27 |
| 6 | 0 | F | 60 |
| 7 | 0 | M | 64 |
| 8 | 0 | F | 59 |
| 9 | 0 | M | 35 |
| 10 | 0 | M | 37 |
| 11 | 0 | M | 68 |
| 12 | 0 | M | 73 |
| 13 | 0 | F | 49 |
| 14 | 0 | M | 83 |
| 15 | 0 | M | 79 |
| 16 | 0 | F | 69 |
| **group medians (IQR)** | | **50 : 50** | **66 (21.75)** |
| 17 | I | F | 70 |
| 18 | II | M | 87 |
| 19 | II | M | 52 |
| 20 | III | M | 79 |
| 21 | III | M | 90 |
| 22 | II | M | 70 |
| 23 | II | F | 66 |
| 24 | III | F | 77 |
| 25 | II | F | 86 |
| 26 | III | F | 91 |
| 27 | II | F | 61 |
| 28 | II | M | 83 |
| 29 | III | F | 85 |
| 30 | I | M | 66 |
| 31 | II | M | 80 |
| 32 | III | M | 88 |
| **group medians (IQR)** | | **43.75 : 56.25** | **79.5 (17.25)** |
| 33 | VI | F | 69 |
| 34 | VI | M | 78 |
| 35 | V | F | 96 |
| 36 | VI | F | 75 |
| 37 | VI | F | 64 |
| 38 | V | M | 82 |
| 39 | V | M | 89 |
| 40 | VI | F | 79 |
| 41 | V | M | 71 |
| 42 | VI | M | 66 |
| 43 | VI | F | 78 |
| 44 | V | M | 88 |
| 45 | VI | F | 74 |
| 46 | IV | M | 86 |
| 47 | VI | F | 81 |
| 48 | V | M | 88 |
